# Live Spike Sorting of Large-scale Neural Recordings

**DOI:** 10.64898/2025.12.29.696938

**Authors:** Shreyas Muralidharan, Calvin Leng, Lucas Orts, Ethan Trepka, Shude Zhu, Matthew Panichello, Donatas Jonikaitis, Frank Lanfranchi, Jacob Pennington, Marius Pachitariu, Tirin Moore

## Abstract

Online monitoring of neuronal activity has tremendous value for closed-loop control in both experimental and clinical settings, yet established methods rely primarily on thresholded neural activity, rather than sorted single-neuron spikes. The recent introduction of large-scale electrophysiological tools has greatly expanded single-neuron recording capacity in animals and humans highlighting the need to sort spikes during data collection. Here, we describe a system for live spike sorting (LSS) populations of hundreds of neurons with millisecond-scale latency. Using neurophysiological recordings from macaque visual cortex with Neuropixels probes, we show that LSS closely replicates the temporal responses and tuning of single neurons obtained using offline sorting. We further show that decoding neural signals with LSS achieves the same performance as that obtained from offline sorting. Lastly, we demonstrate the capacity of LSS to enable closed-loop interventions based on the activity of specific neuronal subclasses.

## Introduction

The use of online measurements of neural activity has played an important role in both basic and translational neuroscience. In basic neuroscience, online measurements of different levels of neural activity (e.g. membrane voltage, spiking, EEG) have been used to introduce closed-loop perturbations to address specific experimental hypotheses that directly test the contributions of neural states to behavior (e.g. ^1–6^). By delivering interventions on the timescale of neural processing, such approaches allow experimenters to modulate ongoing computations and directly observe their behavioral consequences. For example, previous closed-loop experiments involving neuronal spiking activity have been used to demonstrate the role of prefrontal activity in selective attention^7^, the influence of decision variables on choice behavior^8^, and to discover the optimal stimuli for visual cortical neurons^9^. Notably, measurements of spiking activity in these and other closed-loop studies were limited to small numbers of single neurons or to multi-unit or thresholded neural activity.

In clinical settings, closed loop paradigms make it possible to restore functions to individuals with neurological impairments by harnessing neural signals in real-time to control other devices via brain-computer interfaces (BCIs). BCIs have been used to restore functions in individuals with motor or communication impairments via online decoding of handwriting^10^, reaching and grasping^11^, and speech^12,13^, and to optimize parameters in therapeutic neuromodulation^14^. Achieving optimal performance requires reliable, real-time extraction of neural activity from large neural ensembles, requiring decoding algorithms capable of operating on millisecond timescales. However, at present, spiking-based BCIs rely primarily on thresholded neural activity rather than isolated single-neuron spikes, thus obscuring information available from different subclasses of neurons.

The recent advent of large-scale, high-density electrophysiology has enabled the recording from hundreds of neurons simultaneously. This approach uses multi-electrode arrays (MEAs) with large channel counts (100s) and spatially dense electrode layouts (e.g. 20 µm spacing) that support high-quality spike sorting of extracellular single-neuron activity. A noteworthy example of MEAs is the recently developed Neuropixels probes. Neuropixels are silicon-based probes with continuous, dense, and programmably selectable electrode contacts. To date, they have been used widely in a variety of model species^15–23^ including nonhuman primates^24–26^ and humans^27,28^.

The proliferation in the use of MEAs highlights the need to provide a system for achieving high-quality spike sorting online that is easy to implement and well-integrated with widely used large-scale neurophysiological tools. A key limitation of the standard neurophysiological workflow is that it precludes experimental interventions based on the activity of large local or distributed populations of single neurons. A wealth of studies have linked spatiotemporal patterns of neuronal population activity to fundamental components of perception, cognition and behavior ^17,24,26,29–32^. Further evidence suggests specific roles of electrophysiologically distinguishable neuronal subclasses (e.g. ‘fast-spiking’ neurons) in those functions^33–37^, thus indicating a need for a system to measure signals from neuronal subpopulations online. Such a system could naturally be deployed in BCI settings as well. To address this gap, we developed a system for online, “live”, spike sorting (LSS) that enables access to single-neuron activity at short latencies. Our system builds on the widely used Kilosort platform^38,39^ to facilitate adoption by the broad community of neurophysiologists.

## RESULTS

### LSS Implementation

To date, a number of algorithms have been developed to transform extracellular multi-electrode recording data into single-neuron clusters, recovering each neuron’s spatiotemporal electrical footprint across channels and its corresponding spike times. At present, the vast majority of laboratories utilizing MEAs, particularly Neuropixels probes, use Kilosort for spike sorting^24–26,28,38,40–43^. We chose to develop LSS on the Kilosort platform for this reason, though it could also be integrated with other sorting platforms. Typical MEA workflows follow a two-stage pipeline (Figure 1, top). During the recording phase, extracellular electrophysiological data is acquired and written to disk using standard data acquisition systems and software (e.g. SpikeGLX). In the second stage, offline spike sorting is performed using the chosen sorting algorithm, producing estimates of neuron locations, spike templates, and spike times. These outputs are then analyzed in relation to experimental variables to study neural circuits. LSS establishes a new workflow in which a short (e.g. 10-15 minutes) training period of recording is first used to establish waveform spiking templates (Figure 1, bottom). To determine a sufficient training period duration, we examined the duration and characteristics of recordings needed to obtain reliable waveform templates (Supplementary Information). Next, subsequently acquired raw electrophysiological data are processed and spike sorted live, using the clusters established in the training period. For our purposes, we define live as the ability to sort activity from short intervals (e.g. 50 ms) of data in less than the duration of that interval. From this, activity can then be monitored in the graphical user interface (Supplementary Video) and analyzed on the timescale of changing experimental variables.

**Figure 1.**
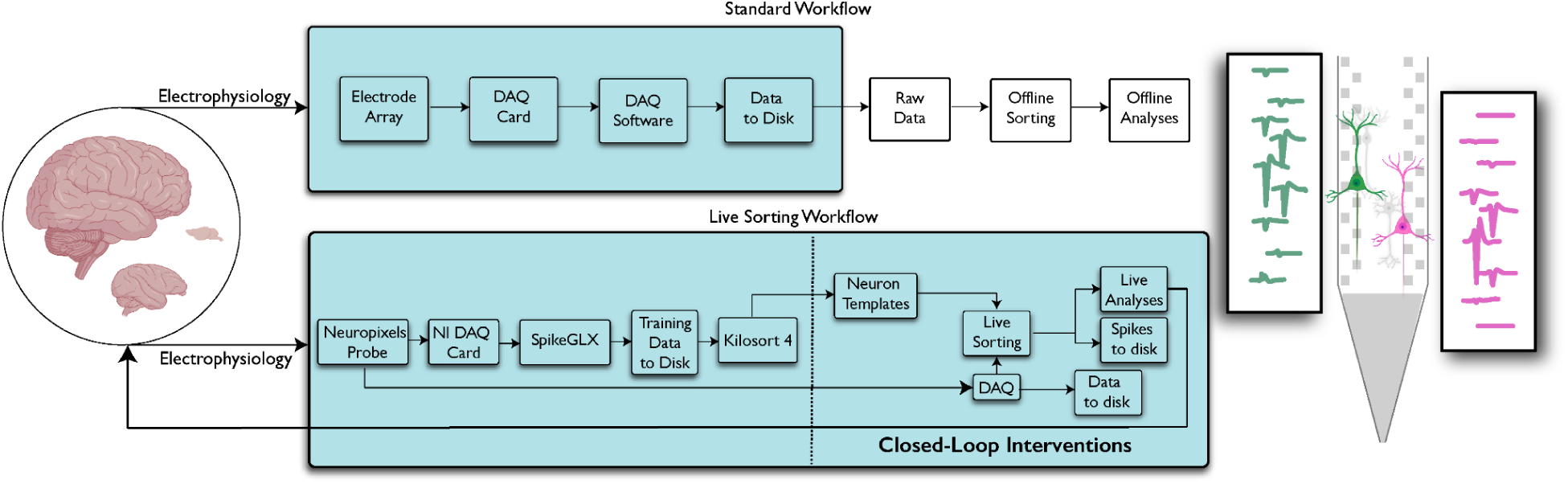
Standard and live spike sorting workflows. Top, in the standard workflow, data are acquired during a recording session (shown in teal), and written to disk. Subsequently, stored experimental data are accessed for offline sorting (e.g. Kilosort4) to retrieve single-neuron information (waveform locations, templates, and spike times) which are then used for analyses. Bottom, in the live spike sorting (LSS) workflow, a two-step process (separated by dotted line) begins with an initial training recording. Sorting is then performed on the training data. Templates and neuron information from these data are then used to sort subsequent data live, while simultaneously writing the same data to disk. Spike data sorted live can then be analyzed during the recording, e.g. for closed-loop interventions. The right cartoon depicts multi-electrode recordings from adjacent neurons and the post processed waveform templates across locations.

### LSS Benchmarking

We benchmarked the performance of LSS in sorting spikes live using Neuropixels recordings obtained from large populations of visually responsive neurons in macaque visual cortex. We chose visual cortex as a testing ground for LSS given that neurons there exhibit relatively reliable, interpretable, low-latency and distinctly selective responses to sensory stimulation which can be used to exemplify the benefits of single-neuron isolation. Neurons were recorded in the superior temporal sulcus (STS), primarily area MT, while monkeys performed a standard fixation task (Methods). Using the uncurated Kilosort4 output, these sessions yielded a total of 6,986 isolated single and multi-neuronal clusters (419-957 neurons/session).

To evaluate the performance of LSS, we compared its readout of spiking activity after the training period to that obtained from the offline workflow. We compared LSS to the offline output in three ways. First, to assess whether LSS could reproduce the fine temporal spiking dynamics of individual neurons, we compared the peri-stimulus time histograms of matched units clustered via the two methods. Second, we assessed the similarity of visual motion selectivity output by LSS with that of offline by comparing the tuning curves of matched units. To perform the above two comparisons, units from the LSS training period were matched to units obtained offline (Methods). From this, we obtained 1,029 matched neurons across the ten sessions. Third, we compared the performance of decoders trained on the total population activity retrieved offline or live with LSS.

#### Single unit peri-stimulus time histograms (PSTHs)

We first compared the spike trains of single units retrieved live with LSS to those retrieved offline using the full set of matched units. During each recording in area MT, drifting grating stimuli were presented multiple times at the location of neuronal RFs thus evoking multiple responses from neurons during each trial (Figure 2a). This provided an opportunity to assess similarities in the spike trains of individual unit responses retrieved live and offline. Thus, we compared the PSTHs generated from the two methods. We found that the PSTHs were highly correlated across matched units (median Pearson correlation = 0.96, IQR = 0.18) (Figure 2b). This demonstrates that LSS recapitulated the fine temporal dynamics of visual cortical responses obtained by offline spike sorting.

**Figure 2.**
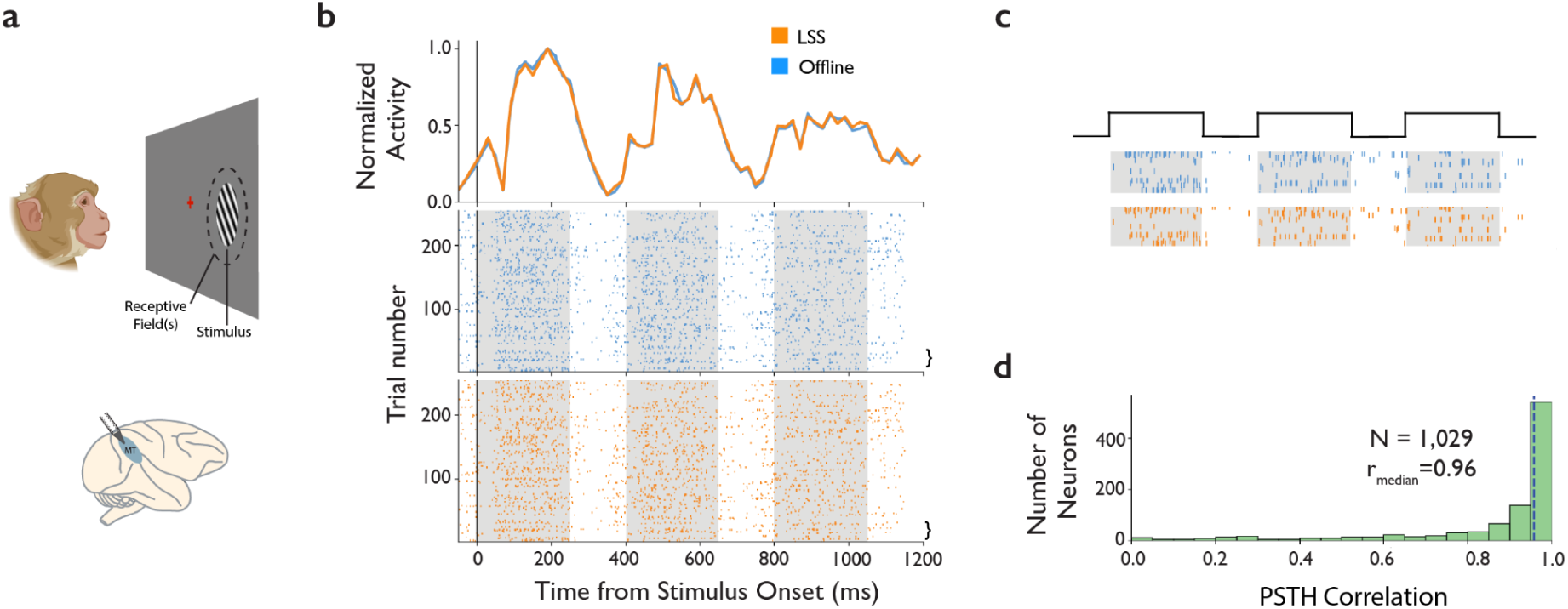
Comparison of live and offline sorted peri-stimulus time histograms. A, monkeys performed a fixation task and the activity of neurons in area MT (top) was measured in response to drifting gratings varying in direction. B, Response of an example area MT unit, aligned to the onset of the first of three visual stimulus presentations in each trial. Top, peri-stimulus time histograms are shown for activity retrieved with LSS (orange) and offline (blue). Bin size, 100 ms; step size, 10 ms. Bottom, raster plots of live and offline activity. Gray shading denotes the onset and duration of stimulus presentations. C, Magnified view of spike rasters from several example trials (brackets) in B. D, Distribution of correlations between peri-stimulus time histograms for all live and offline sorted units.

#### Stimulus tuning of single neurons

Next, we compared the responses of single units to varying directions of visual motion using activity retrieved live and offline. Area MT is organized into cortical columns in which neurons within the same column exhibit similar preferences to the direction of visual motion^44^. Our dataset consists of recordings made tangential to the cortical surface and consequently pairs of neurons located further apart tended to exhibit larger differences in their preferred motion direction (Figure 3a). For each of the 1,029 matched units, we fit von Mises functions to their responses across motion direction. Of these, 309 units were well fit (r^2^>0.6) in both the live and offline retrieved activity. We found that the tuning peaks of these units were very similar between live and offline methods (Figure 3b) (mean absolute difference = 10.75 deg.). Lastly, to measure the tuning similarity for all matched units, we computed Pearson correlations between tuning curves derived from LSS and offline sorting. Overall, tuning curves were highly correlated between the two methods (median = 0.91, IQR = 0.32) (Figure 3c). Thus, in addition to yielding similar spike trains, LSS also recapitulated the tuning curves of single units across stimulus conditions.

**Figure 3.**
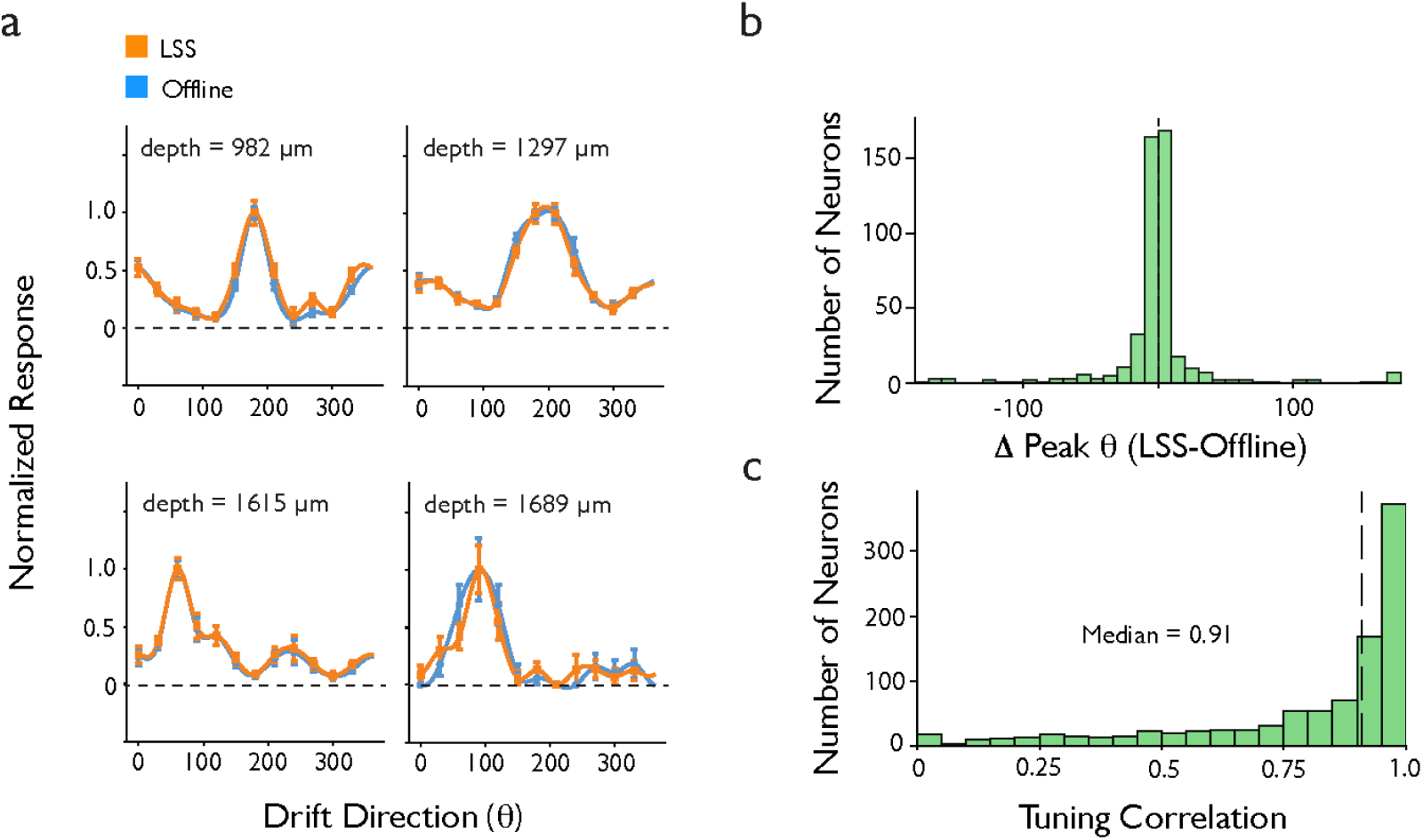
Comparison of live and offline sorted tuning curves of area MT neurons. A, tuning to the direction of visual motion of drifting gratings from four example area MT neurons in a single recording session, LSS (orange) or sorted offline (blue). Responses of each neuron were normalized to the peak firing rate and are shown with spline interpolation. Error bars denote SEM. Recording depths are shown for each neuron from shallow to deep. B, Distribution of differences between online and offline tuning peaks for neuronal responses well fit by the von Mises function (n=309). C, Distribution of correlations between tuning curves for all live and offline sorted units (n=1029).

#### Stimulus decoding from populations of neurons

A major advantage of sorting single-unit data live is that the sorted activity can then be used to decode signals conveyed by populations of individual neurons. Thus, to further validate LSS, we measured the extent to which population decoding from activity sorted live achieved comparable performance to offline decoding. Using data from each of the recording sessions from area MT, we trained multi-class logistic regression classifiers separately on spiking activity retrieved live with LSS and offline (Methods). Each classifier was trained to decode one of twelve directions of grating drift across 100 ms time bins around the time of stimulus onset.

We first compared the performance of decoders trained on all units (features) available from each session’s data using offline and live sorted activity. Across the 10 sessions, as expected, classifiers trained on offline sorted activity robustly decoded the direction of motion following stimulus onset (72.3% +/- 2.3%; chance performance = 8.33% , 50-500 ms) (Figure 4a) (Extended Data Figure 1). The performance obtained from classifiers trained on live sorted activity using LSS was comparable to that of offline classifiers (70.0% +/- 2.9%), with an average reduction of 2.2%. This was true despite offline sorting consistently identifying more units than live sorting (Offline n= 6,986; LSS n=3,956 across sessions), due to the greater duration of offline sorted data. To compare decoding performance independent of this difference, we computed feature dropping curves^45^ from each session’s data for offline and live sorted activity. These curves were then fitted with asymptotic functions to extract the number of features at which performance saturated (asymptotic performance) (Figure 4c) (Extended Data Figure 2) (Methods). Across the 10 sessions, the performance of decoders trained on offline sorted data required more units to achieve asymptotic performance compared to live sorted data (mean difference 91.4 units; p<0.05). This is consistent with the greater unit yield obtained with offline sorting. Critically, however, the asymptotic performance yielded from live sorted data was statistically indistinguishable from that obtained from offline sorting (mean difference in performance +/- 1.3%, p>0.05)(Figure 4c). Thus, when controlling for the number of units, LSS achieved the same decoding performance as that obtained with offline sorting. In addition, we found that neither the probe drift rate (r=-0.043, p>0.05) nor the duration of the test recordings (r = -0.385, p>0.05) significantly correlated with differences in performance between LSS and offline sorting (Extended Data Figure 3).

**Figure 4.**
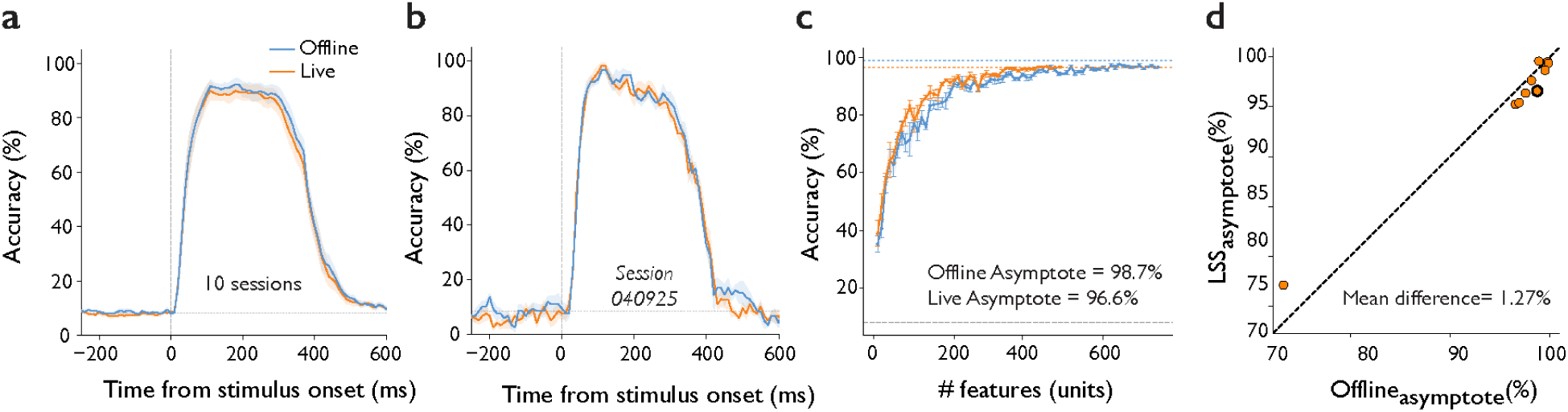
Comparison of population decoding from live and offline sorted activity. A, Average classifier performance time course for 12-way visual motion direction decoding across populations of area MT neurons, averaged over 10 recording sessions in two monkeys. Traces compare offline sorted spikes and live sorted spikes (LSS), aligned to stimulus onset. Gray dashed line denotes chance performance (8.3%). B, Classifier performance time course for a single example session. C, Feature-dropping curves for live sorted and offline sorted activity in the session shown in B. Dotted horizontal lines indicate fitted asymptotic accuracy for each method. Dashed line denotes chance; error bars denote SEM. D, Comparison of asymptotic decoding accuracy between live sorting and offline sorting across all 10 sessions. Each point represents one session; the highlighted point corresponds to the example session in B and C. Dashed line denotes unity. The mean difference in asymptotic accuracy was 1.27 % (Wilcoxon signed-rank test, p >0.05).

### Use of LSS in closed-loop experiments

Lastly, we sought to demonstrate the utility of LSS in closed-loop neurophysiological experiments. Closed-loop experiments enable investigators to test the role of neural states in brain function^1–7^. To achieve this, we live sorted neuronal activity recorded in area MT as a means of observing the influence of endogenous neural states on sensory responses. It is well known that sensory neurons in neocortex exhibit spontaneous activity, and the relationship of that activity to sensory driven activity has been the focus of numerous studies ^46–50^. Although general principles governing the influence of spontaneous activity on sensory coding in the cortex remain elusive, evidence of such an influence is clear. For example, Gutnisky et al. demonstrated that the state of pre-stimulus activity influences the visually evoked responses of macaque V1 neurons. In this case, investigators compared the visual responses occurring on trials in which pre-stimulus single-neuron firing rates (FRs) were either high or low, based on offline measurements. In such a design, the pre-stimulus FR is not under experimental control and the difference between high and low states is thus limited by the coincidence of stimulus onset with spontaneously fluctuating activity. We used LSS to maximize the difference between high and low neural states by controlling the timing of stimulus presentation with neuronal activity.

We used LSS to trigger stimulus presentations based on the activity of populations of live-sorted MT neurons (Methods). Specifically, we triggered stimulus presentations on the activity of ‘fast-spiking’ (FS) neurons (Figure 5a). Past neurophysiological studies have established that different morphological subclasses of neocortical neurons can be distinguished to some extent by their extracellularly recorded waveforms ^51–53^. Most often, FS waveforms are thought to identify putative inhibitory neurons, whereas ‘regular-spiking’ (RS) neurons are thought to identify excitatory neurons ^53–55^, though this distinction has important exceptions ^56–58^. Our focus on the activity of FS neurons thus allowed us to trigger stimulus events on the activity of a subclass of neurons likely to have functionally distinct properties. During the experiment, on each trial, a drifting grating was presented in the neuronal RFs while the monkey maintained central fixation. Stimulus presentation was triggered either when the mean FR of the FS population exceeded a threshold value (Figure 5a), or at the end of a 1-second period (“non-triggered trials”).

**Figure 5.**
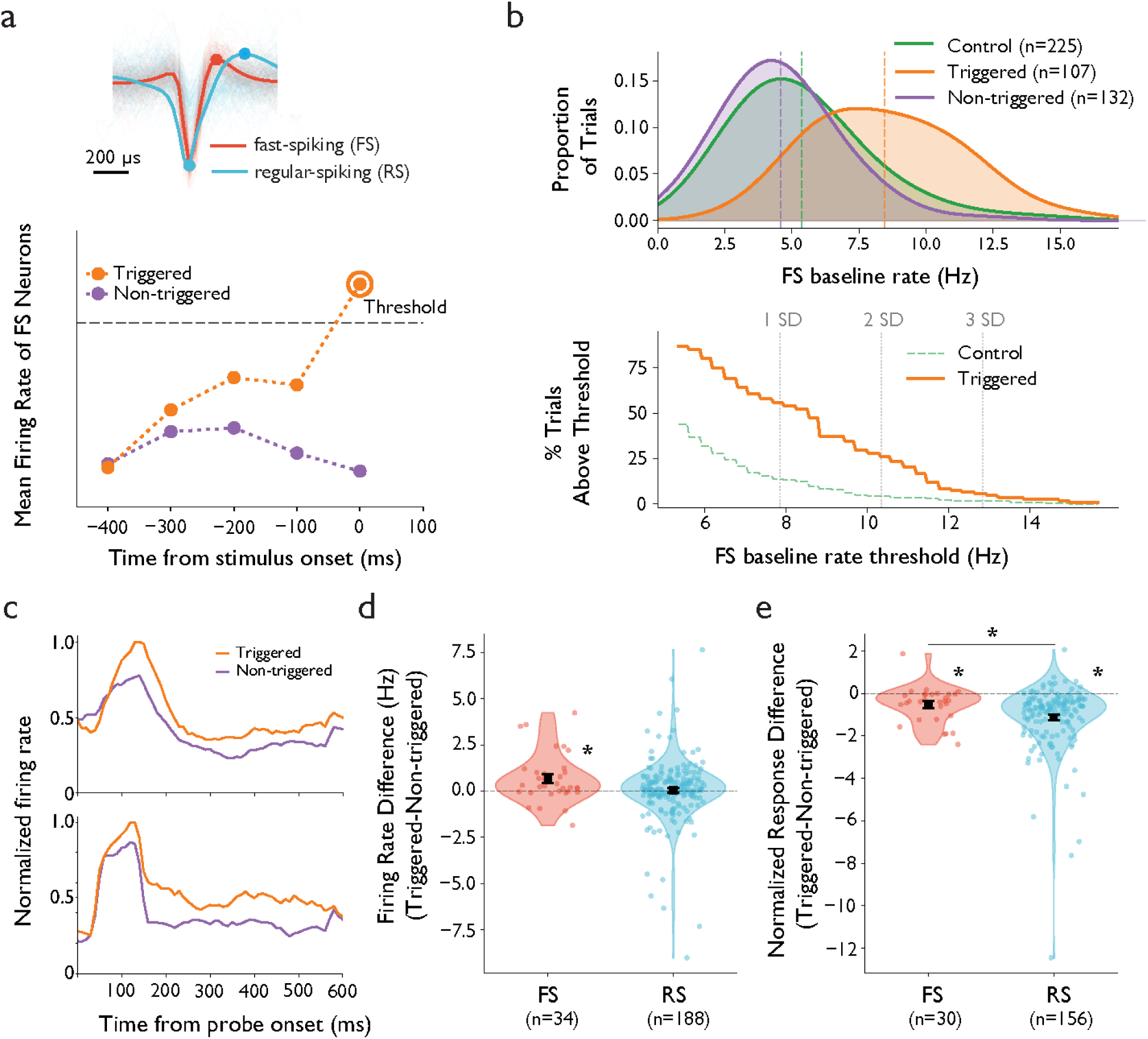
Closed-loop triggering of visual stimulation with LSS. A, top, example spike waveforms of an example fast-spiking (red) and regular-spiking (blue) neuron. Filled circles denote trough and peaks for both waveform classes. Bottom, schematic of the closed-loop experiment depicting the onset of stimulus presentation triggered by an above-threshold firing rate of FS neurons (triggered), or at the end of the fixation period (non-triggered). Plot shows the time course of FS firing rates on triggered (orange) or non-triggered trials (purple). The circled point on the triggered trial denotes the threshold time bin. B. Top, Distributions of firing rates of fast-spiking neurons during control (green), non-triggered (purple), and triggered (orange) trials. Dotted lines indicate the mean of each distribution. Bottom, percentage of trials above a given threshold for triggered trial (orange) and in control trial distributions (green) C. Single neuron PSTHs for two example FS neurons, shown for triggered and non-triggered trials. D. Left, distribution of differences in visually evoked firing rates (triggered - non-triggered) for FS (red) and RS (blue) neurons. Right, differences in visually evoked responses normalized (divided) by pre-stimulus activity. Only units with non-zero pre-stimulus activity are included. Asterisks denote significance at p<0.01.

After offline sorting of neuronal activity from the closed-loop experiment, we compared the distributions of pre-stimulus (−100 to 0 ms) FRs of FS neurons (n=34) across triggered trials, non-triggered trials, and control trials with conventional stimulus presentation. Control trials were used to obtain an unbiased estimate of the pre-stimulus FR distribution. This comparison revealed significant differences in FRs between all 3 subsets of trials (control mean_pre-stimulus_=5.4 Hz; non-triggered mean_pre-stimulus_=4.6 Hz; triggered mean_pre-stimulus_=8.5 Hz; p<0.01) (Figure 5b). Notably, the percentage of triggered trials above a given FR threshold was much greater than the percentage that would have been obtained on control trials. Thus, triggering substantially increased the difference between high and low FR states for a given trial count.

Finally, we compared the visually evoked responses between triggered and non-triggered trials. Consistent with previous evidence of baseline-dependent sensory responses^49,59^, we observed that the visual responses of FS neurons evoked during triggered trials were greater than those evoked on non-triggered trials (Figure 5c). This effect was significant across the population of fast spiking neurons (triggered mean_response_ = 8.0 Hz; non-triggered mean_response_=7.4 Hz; p<0.01) (Figure 5d). In contrast to FS neurons, the pre-stimulus effect on visual responses was absent in the larger population of RS neurons (n=188). That is, although the pre-stimulus FR of RS neurons was greater on triggered trials than non-triggered trials (non-triggered mean_pre-stim_=3.5 Hz; triggered mean_pre-stim_=6.9 Hz; p<0.01), their visual responses were not (triggered mean_response_ = 7.8 Hz; non-triggered mean_response_=7.8 Hz; difference, p=0.7).

In addition, to measure visual responses independent of pre-stimulus firing rates, we normalized each neuron’s visual response to its pre-stimulus FR. This revealed a significant reduction in visual responses in both FS neurons (triggered median_response_ = 0.92; non-triggered median_response_ = 1.1; difference, p<0.05) and RS neurons (triggered median_response_ = 1.1; non-triggered median_response_ = 2.0; p<0.0001) (Figure 5d) , a result also consistent with previous studies^49,59^. Moreover, this reduction was greater in the RS neuron population than the FS population (Mann-Whitney U test, p<0.005). Thus, for both normalized and non-normalized measures, the effect of pre-stimulus FR on visually evoked activity differed between the two putative cell types. Future experiments utilizing live spike sorting could further explore the role of specific cell types in shaping the relationship between spontaneous and stimulus-driven activity. For example, in addition to triggering instead on RS activity, stimulus presentation could also be triggered on contrasting FS and RS FRs as a means of probing an influence of excitatory/inhibitory balance on stimulus coding ^60^. More generally, LSS enables a shift from post hoc analyses of neural states to real-time experimental control based on the activity of identified neurons, thus providing a tool to investigate how specific cell types and population dynamics contribute to perception, cognition, and behavior.

## DISCUSSION

Here, we have described LSS, a system for sorting neuronal spiking activity as it is collected live from high-density microelectrode probes. The system leverages the technical advantages and broad use of Kilosort4 to provide a robust and easily implementable system for obtaining online analyses of single-neuron spiking data. Live sorting with LSS is achieved by utilizing an initial period of recorded neural data to identify waveform templates and then assigning incoming data to trained templates. We benchmarked LSS by comparing its output to that obtained with offline sorting. We show that LSS closely replicates the peri-stimulus time histograms and visual response tuning curves of single-neurons obtained using offline sorting. We also show that decoding neuronal signals online with LSS achieves equivalent performance as that obtained with offline sorting. Lastly, we demonstrate the utility of LSS in a closed-loop experiment in which visual stimulation was triggered based on the spontaneous activity of a subclass of neurons.

As this is the first iteration of LSS, we have considered a number of ways in which the system might be advanced further to improve upon its performance and robust alignment with the quality of offline sorted data. First, estimation and correction of recording drift could be done live using principles from the Kilosort4 algorithm, allowing for improved tracking of clusters identified in the training period. Furthermore, simultaneous updating of training clusters could be achieved by running offline Kilosort on data written to disk during live sorting. Thus, the training templates could be updated to enable live sorting of units that may become detectable after the conclusion of the training period. Second, the addition of manual curation of units found in the training period could easily be achieved to impose stricter criteria on live sorting. Finally, integration with other data acquisition platforms such as Open Ephys could enable more widespread adoption of LSS.

As a proof of concept, we demonstrated the benefit of LSS in a closed-loop neurophysiological experiment by live-sorting neural activity and using it to control the timing of visual stimulus presentation. Specifically, stimulus onset was triggered by the state of pre-stimulus FR of fast-spiking neurons in area MT thus enabling us to substantially enrich the proportion of trials occurring during high FR states relative to what would be obtained by chance. This approach addresses a limitation of conventional experimental designs, in which the sampling of particular neural states is constrained by their spontaneous occurrence. A wealth of evidence suggests that many fundamental brain functions, e.g. perception ^30,61–63^, working-memory^24^, decision-making ^8,64,65^, and motor preparation ^66–68^, are associated with unique, temporally discrete patterns of neuronal activity^69^. When these patterns of activity are rare and/or transient, conventional designs require large numbers of trials to accumulate sufficient observations for post hoc analyses. Our results demonstrate the utility of live spike sorting in enabling investigators to identify targeted patterns of single-neuron activity as they occur in closed-loop experiments, and more directly investigate their role in brain function.

The closed-loop framework enabled by LSS extends naturally to a range of experimental manipulations beyond those demonstrated here. For instance, rather than triggering a single stimulus class, investigators could use neural state detection to select among multiple stimulus conditions, enabling within-session comparisons of how different stimuli interact with the same neural state. Similarly, LSS could be used to trigger direct perturbations of neural activity, such as optogenetic or electrical stimulation, contingent on the activity of specific cell types. Real-time decoding of population activity with LSS could also be used to trigger behavioral task events on latent neural states^70,71^, rather than simply on FR, enabling more sophisticated interrogation of the relationship between neural coding and behavior. More broadly, because LSS provides continuous access to single-neuron identity during recording, it allows for the possibility of designing adaptive experiments in which trial parameters are iteratively updated based on accumulating neural data, an approach that could substantially accelerate the characterization of high-dimensional neural response subspaces^72,73^.

## Supporting information

Supplemental Figure and Description

## Acknowledgements

The authors would like to thank Danielle Lopes and Stephen Cital for veterinary assistance. This work was supported by NIH grants EY014924 and NS11662302, and a Ben Barres Professorship to T.M..

## METHODS

### LSS Implementation

#### System architecture

The system interfaces with SpikeGLX to receive continuous multichannel extracellular recordings at 30 kHz. Data are acquired in variable-length batches and transferred to GPU memory via pinned host buffers. Adjacent batches overlap by twice the template duration (2M samples) to prevent spike loss at boundaries. When processing cannot keep pace with acquisition, an adaptive skip strategy discards older samples to ensure the most recent activity is always processed. Multiple GPUs can operate in parallel on separate probes or channel subsets.

#### GPU Implementation

The original Kilosort4 pipeline uses PyTorch for GPU acceleration in an offline batch-processing context. Our system reimplements the computationally intensive stages directly in CUDA C++ to eliminate Python and framework overhead and to maintain deterministic, low-latency execution suitable for real-time operation. The two dominant computational steps (PCA projection and template cross-correlation) are expressed as large matrix multiplications dispatched through cuBLAS, which saturates GPU throughput on modern hardware. High-pass filtering is performed via cuFFT-based convolution rather than a recursive Butterworth filter, enabling the entire filter operation to execute as a single GPU kernel launch. Custom CUDA kernels handle the remaining operations: 1D max-pooling for local maxima detection, spike amplitude computation, PCA feature extraction at neighboring channels, and mass-weighted position estimation. These kernels are designed to minimize memory round-trips by operating in-place on device memory wherever possible. Host-device data transfer uses pinned (page-locked) memory, which enables DMA transfers that proceed concurrently with GPU computation on successive batches. All intermediate buffers (residuals, convolution results, PCA projections) remain in GPU global memory throughout the matching pursuit loop, avoiding repeated transfers across the PCIe bus. The result is that the only CPU-GPU transfers per batch are the initial raw data upload and the final spike index/amplitude download.

#### Offline template learning

Prior to online operation, Kilosort4 is run on an initial recording segment to learn spike templates and preprocessing parameters. The live system loads the resulting template waveforms, PCA basis vectors, PCA-space template features (Wall3), pairwise template cross-correlations, spatial whitening and drift correction matrices, high-pass filter coefficients, channel geometry, and PCA-space cluster centroids.

#### Online preprocessing

Each batch undergoes GPU-accelerated preprocessing: (1) per-channel mean subtraction, (2) common median referencing across channels, (3) FFT-based high-pass filtering at 300 Hz, (4) spatial whitening via the precomputed whitening matrix, and (5) drift correction via the precomputed drift matrix.

#### Template matching via matching pursuit

Spike detection follows the matching pursuit framework of Kilosort4, reimplemented in CUDA. The preprocessed batch is first projected into PCA space and cross-correlated with all template features via cuBLAS matrix multiplication. An iterative detection loop (up to 50 passes) then: normalizes the convolution output (ReLU, squaring, border zeroing), identifies local maxima via max-pooling in time, thresholds detections at a learned threshold, computes spike amplitudes, and subtracts each detected spike’s contribution from both the residual signal and the convolution result using precomputed template cross-correlations. Iteration terminates when no spikes exceed the threshold.

#### Spike localization and cluster assignment

Detected spikes are localized on the probe by extracting PCA features from the residual at nearby channels and computing a mass-weighted average of channel positions. Pre-clustered template identities are mapped to final neuron clusters via nearest-neighbor assignment in PCA feature space using centroids from the offline Kilosort4 run. Duplicate spikes are suppressed by enforcing a minimum inter-spike interval per template.

### Setting up LSS

LSS can be used with any CUDA enabled GPU. We tested the system with CUDA version 11.3. LSS can be found at this link: https://github.com/MShreyasStanford/LiveSpikeSorter, and we recommend using the Microsoft visual studio compiler and linker (cl.exe). We have not tested compilation of this project under any standard other than C++17. Components of the system, such as the LSS or Python GUI can be modified to fit specific use cases. Detailed instructions can be found at the github.

### Neuron Matching-LSS

To evaluate LSS performance on real recordings, we trained LSS on the first 15 minutes of each recording (the training period) and generated spike trains for the full recording duration. As a reference, we ran Kilosort4 offline on the full recording. Because LSS template initialization and clustering are derived from the training period, unit identities are reliably defined there; we therefore restricted unit matching between LSS and offline Kilosort4 to spike trains from the 15-minute training period. We determined unit matches as in the Kilosort4 paper^38^. We computed false positive rate (FP) and false miss rate (FM) for pairs of spike trains and defined their match score as 1-FP-FM. Units with a score of at least 0.8 are considered matched.

### Electrophysiological recordings

#### Experimental subjects

Neural recordings were collected in a total of ten sessions from two male Rhesus macaques (A: age 14 years, weight 11 kg, H: age 13 years, weight 17kg). All surgical and experimental procedures were approved by the Stanford University Institutional Animal Care and Use Committee and were in accordance with the policies and procedures of the National Institutes of Health.

#### Electrophysiology

We recorded spiking activity from neurons using primate Neuropixels probes (IMEC). In each recording session, the dura was perforated with a 21-gauge pointed cannula mounted on an independent micromanipulator. A single Neuropixels NHP probe was then advanced through the cannula using either a hydraulic Narishge microdrive or a motorized drive system. Recording sites were configured as 384 simultaneously active channels in a contiguous block (3.84mm span), providing dense sampling along the probe shank. Neuropixels probes digitized signals at the headstage and streamed separate bands for action potentials (AP; 300 Hz high-pass; 30 kHz sampling) and local field potential (LFP; 1 kHz low-pass; 2.5 kHz sampling). Neural activity was monitored online and saved to disk using SpikeGLX (https://billkarsh.github.io/SpikeGLX/).

#### Behavioral task and recording setup

Neural recordings were conducted in the superior temporal sulcus (STS) while monkeys performed a fixation task in which visual stimuli were presented to map receptive fields (RFs) and to measure direction of motion selectivity. The location of the STS was identified using anatomical MRIs in both monkeys and confirmed based on the functional properties of recorded neurons including RF size and robust selectivity to planar visual motion which indicated that most neurons were in area MT. Neural data were recorded with Neuropixels 1.0 NHP probes.

Experiment code was written in MATLAB (MathWorks, version R2020b) using the Psychophysics Toolbox^74^. Stimuli were presented on a ViewPixx LCD monitor with 1920×1080 pixels resolution and 100 Hz refresh rate (VPixx Technologies). The monkeys viewed the display from 42 cm. Eye position was monitored at 1000 Hz using monocular corneal reflection and pupil tracking with an Eyelink 1000 Plus (SR Research Ltd., Ottawa, ON, Canada). Eye-tracker calibration was performed with a five point protocol at the beginning of each recording session.

#### Visual Stimuli

The stimuli used to measure direction of motion selectivity were circular sinusoidal drifting gratings (10 deg./s, 1 cycle/deg.). Stimuli drifted in one of twelve directions in 30° increments spanning 0-330°. In the task, the monkey fixated a central spot to initiate a trial, then three stimuli were presented in sequence, each for 0.25 s with an interstimulus interval of 0.15 s. Grating stimuli were randomly interleaved with additional plaid stimuli that are not analyzed here. In each recording session, the visual responses to the grating stimuli were measured over a duration of 5-28 minutes.

### Data Analysis

#### Population decoding

To evaluate the quality of neural signals obtained from each spike sorting method, we performed time-resolved 12-way motion direction decoding across 10 recording sessions. Spike times were binned into 100 ms sliding windows stepped every 10 ms, spanning -500 to +1500 ms relative to stimulus onset. At each time bin, we decoded motion direction (12 equally spaced directions, 0-330 degrees in 30 degree increments) from the vector of spike counts across all recorded units using multinomial logistic regression (L-BFGS solver, max 1000 iterations). Decoding was evaluated with 5-fold stratified cross-validation (shuffled, random state = 42). Within each training fold, we applied class-balanced subsampling by randomly downsampling each class to the size of the smallest class to mitigate class imbalance. Features were standardized (zero mean, unit variance) within each fold by fitting on the training set and transforming both training and validation sets. Decoding accuracy was averaged across folds at each time bin, and variability was quantified as the standard error of the mean (SEM) across folds. Chance-level performance was 8.3% (1/12). For across-session summaries, per-session accuracy time courses were averaged across sessions, with SEM computed across sessions.

#### Neuron Dropping Curves

To characterize how decoding accuracy scales with the number of available units, we constructed neuron dropping curves for each session and sorting method. For each session, we averaged firing rates across time bins within a 50-400 ms window following stimulus onset, yielding a single feature vector per unit per trial. We then subsampled units in increments of 10, from 10 units up to the total number of available units. At each unit count, 5 random subsets of units were drawn (without replacement, with deterministic seeding for reproducibility). For each draw, per-unit firing rates were z-scored across trials, and 12-way direction classification was performed using multinomial logistic regression (L-BFGS solver, C = 1.0, max 1000 iterations) with 20-fold stratified cross-validation. Training sets were class-balanced within each fold by subsampling to the smallest class. Accuracy was averaged across folds for each draw, then summarized as the mean and SEM across the 5 draws at each unit count.

#### Asymptotic Performance Fits

To quantify the maximum decoding performance attainable with all available units, we fit a two-term saturating exponential model to each neuron dropping curve. The model was defined as follows:

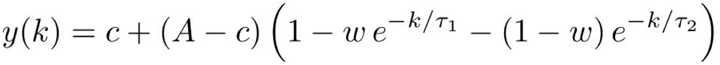

where k is the number of units, A is the asymptotic accuracy, c is chance-level accuracy (1/12, fixed), w is a mixing weight governing the relative contribution of fast and slow rise components, and tau1 and tau2 are fast and slow rise time constants, respectively. The four free parameters (A, w, tau1, tau2) were estimated using nonlinear least-squares fitting (scipy.optimize.curve_fit), with initial guesses set to the observed maximum accuracy, 0.5, 10, and 200 units respectively. Parameter bounds constrained A to [chance, 1.0], w to [0, 1], and tau1 and tau2 to [1e-6, 1e6]. SEM values from the dropping curve were used as inverse weights during fitting. If the fit did not converge, the asymptote was estimated as the mean accuracy of the last 3 points on the dropping curve. Asymptotic accuracy values were compared across sorting methods using a two-sided Wilcoxon signed-rank test on paired (per-session) differences.

#### Statistics

No statistical methods were used to predetermine sample sizes. All statistical tests were two-sided. Pearson correlations were used to assess the similarity of peri-stimulus time histograms and tuning curves between live and offline sorted units. Asymptotic decoding accuracy values were compared across sorting methods using a Wilcoxon signed-rank test on paired per-session differences. Differences in pre-stimulus and visually evoked firing rates across trial conditions (triggered, non-triggered, and control) were assessed using pairwise Mann-Whitney U tests. Differences in normalized visual responses between FS and RS populations were also assessed using a Mann-Whitney U test. All analyses were performed in Python using SciPy (v1.15.2). Significance was defined as p < 0.05 unless otherwise noted.

### Closed-loop experiment

In the example experiment, the LSS training period was derived from a 10 minute recording. In the recording, we targeted a subpopulation of neurons with RFs at a single visual field location using a separate RF mapping procedure (Methods). The subpopulation of neurons corresponded to a subset of channels (260–371) on the Neuropixels probe. Within that population, we used the waveform templates from the Kilosort4 output to classify neurons as either FS or RS based on the trough-to-peak time of each template’s maximum-amplitude waveform (Methods). Neurons with a waveform width less than 200 μs were classified as FS (n =34) ^54^; all others were classified as RS (Figure 5a). The mean and standard deviation (SD) of spontaneous (baseline) activity of the FS population were then computed in 100 ms time windows. Next, we began a set of visual stimulation trials in which a moving grating stimulus was presented while the monkey maintained central fixation. During the fixation period of up to one second, we computed the average FR of the live-sorted FS population every ∼50 ms to detect epochs in which activity exceeded 2 SDs above the baseline mean. Detection of FRs above that threshold immediately triggered the presentation of a moving grating stimulus (triggered trials); otherwise, the stimulus was presented at the end of the fixation period (non-triggered trials) (Figure 5b).

## Notes

### Competing Interest Statement

The authors have declared no competing interest.

### Summary of Updates

This version of the manuscript has been revised to add a figure, correct typos, and revise figures and figure captions.

