## Supplemental Figure and Description for "Live Spike Sorting of Large-scale Neural Recordings"

**Supplemental Information**

*Latencies of LSS steps*

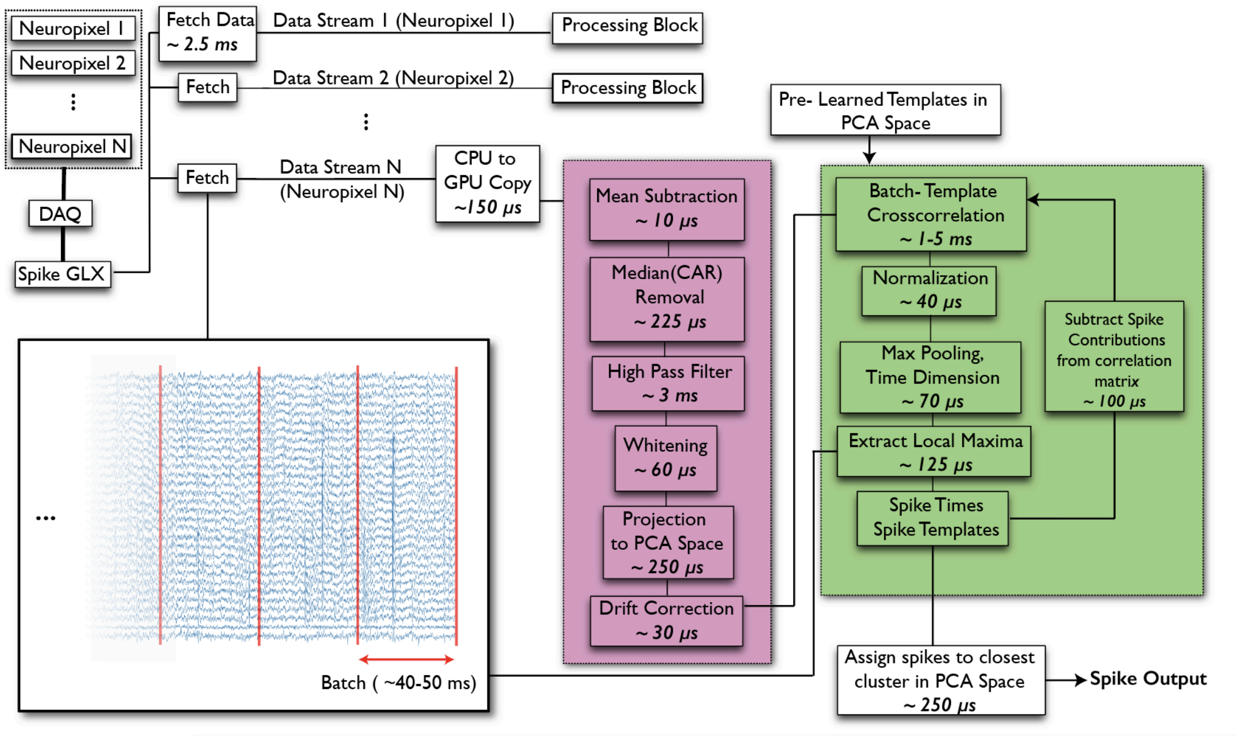


**Supplemental Figure 1: Processing steps for LSS**. Bottom left inset shows a portion of a continuous data stream, processed in small (~40-50 ms) batches (red lines). Each data stream (1-N) is retrieved independently via a fetch call to the SpikeGLX API, followed by a processing block, consisting of pre-processing and spike retrieval. Each batch of data (e.g. N) goes through a series of pre-processing steps (purple) after a CPU to GPU copy. This is followed by spike retrieval via a matching pursuit (MP) algorithm (green). The MP loop terminates when no further spikes remain. The location of each spike in PCA space is used to assign that spike to its closest cluster in the training data.

LSS replicates the key steps of Kilosort4 in an online manner; the mean processing time for each step is shown. In the current version, we assume stability of the recording and assign live sorted spikes to clusters established in the training period based on Euclidean distance in PCA space. This assumption also allowed us to reuse the final drift estimation from Kilosort to pre-process incoming data. While probe drift is an inherent feature of extracellular recordings, a comprehensive characterization of LSS performance across the full range of drift conditions encountered in practice is beyond the scope of this work; we anticipate that the community will further evaluate and refine the system across diverse recording contexts.

We chose to deploy and benchmark the sorter on batch sizes of 50 ms due to the time scale of fluctuation of neural states of interest. However, LSS can be deployed on smaller batch sizes for shorter latencies if needed. We benchmarked the system using both an NVIDIA A6000 GPU and a RTX 3070 GPU. Both GPUs were able to achieve much quicker than live performance while sorting units from the entire 384 channel Neuropixels array, and our results suggest the current implementation could likely process live sorting of units from >384 channels. All code and step-by-step setup instructions are available at: https://github.com/MShreyasStanford/LiveSpikeSorter

*LSS Graphical User Interface*

An important component of the LSS system is the user-friendly graphical interface (GUI) from which the user monitors, selects and evaluates the sorted spike data as it is logged (Supplemental Figure 2). The LSS GUI consists of several panels in which key data are continuously updated, the most basic being the spike rasters of all clustered units. In addition, another key panel continuously updates the accumulating distribution of processing times for all batches. In our current implementation of LSS, each 50-ms batch of data is processed in 9 +/- 3 milliseconds which is largely the sum of times denoted in Supplemental Figure 1. These data can be used to evaluate the extent to which sorting is being performed online. It can also inform hardware decisions and whether sorting should instead be done on a subset of units. Lastly, another panel in the GUI is used to monitor single-neuron statistics for selected units. In the current version, single-neuron statistics such as binned firing rate and sorting quality metrics, e.g. inter-spike interval histograms, and autocorrelograms are included. The supplemental video expands on the GUI.

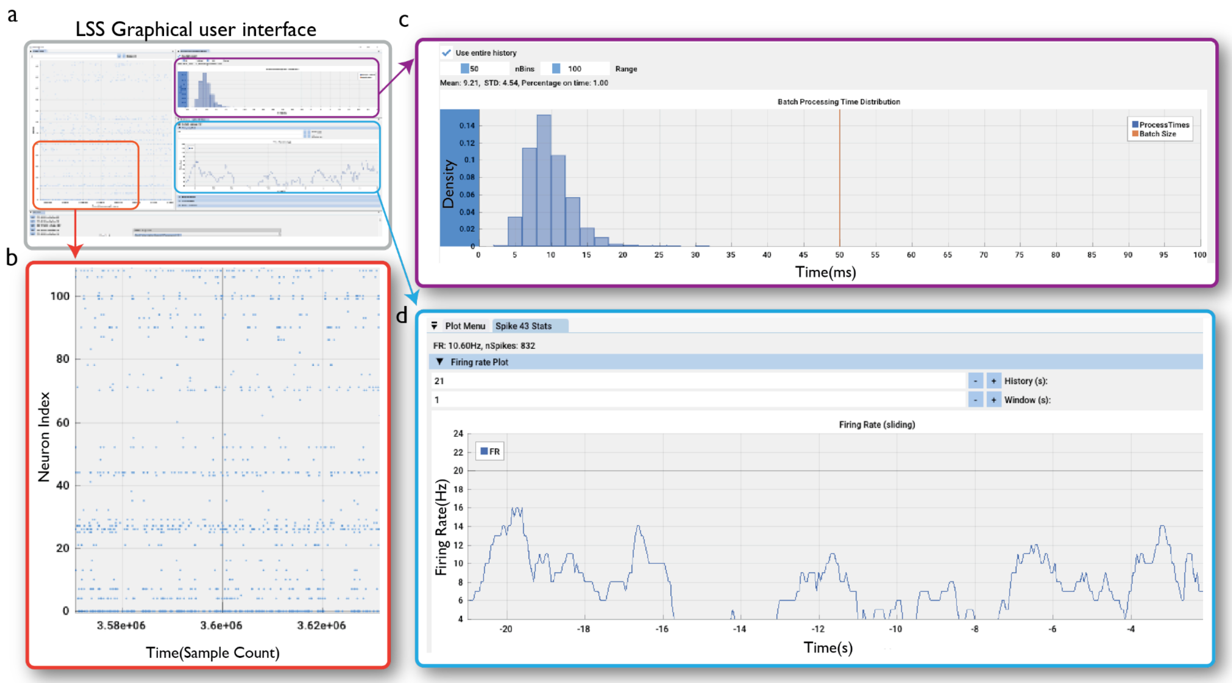


**Supplemental Figure 2. Graphical User Interface for LSS.** A, Live raster of spiking activity across sorted neurons. The x axis displays sample count from the start of SpikeGLX acquisition.

B, histogram of processing times for every 50-ms batch of data, relative to batch duration (orange line). C, Panel to view selected single-neuron statistics, e.g. binned firing rate as shown. Other metrics include spike sorting quality metrics, such as inter-spike interval distributions and autocorrelograms.

**Supplemental Video 1.** Demonstration of the LSS graphical user interface during an active recording session. The video shows real-time updates to the spike raster, batch processing time histogram, and single-neuron statistics as data are acquired and sorted live.

*Validation of LSS Training Period using Kilosort4*

To determine a sufficient duration of training data for the LSS workflow, we first evaluated how well Kilosort4 retrieves the activity of ground truth, simulated units using data of varying durations (Supplemental Figure 3). Simulated datasets with several realistic drift regimes were generated based on real, spike-sorted Neuropixels 1.0 data from the International Brain Laboratory (IBL). Average waveforms were extracted across a large range of depths for sorted units tracked throughout recordings with high drift. Data was simulated by randomly sampling from the collected waveforms at spike times sampled from the ISIs of units in the real data and distributing the waveforms across a virtual probe. Waveform depths were then shifted according to the imposed drift condition. Additional simulation details can be found in Pachitariu et al. 2024.

We determined unit matches as in Pachitariu et al, 2024. Briefly, we computed false positive rate (FP) and false miss rate (FM) for pairs of spike trains and defined their match score as 1-FP-FM. Units with a score of at least 0.8 are considered matched. Only matched units labeled 'good' (Pachitariu et al, 2024) by Kilosort4 with at least a 1 Hz spike rate for the tested portion of the spike trains were considered.  Match rate is defined as the number of matched units divided by the number of units compared. For simulated data, we determined matches against ground truth units. Since different recording lengths were used, we only compared spike trains on the four 5-minute segments shared by all recordings.

Supplemental Figure 3a shows a drift trace along with its estimated drift rates for an example simulated recording with realistic drift. We estimated drift rate from the per-batch drift amounts used for drift correction by Kilosort4. We smoothed these drift amounts with a 2nd-order Savitzky-Golay filter with a window size of 361 batches (12 minutes) to reduce noise, then summed across timepoints the velocity values (first-order absolute differences between consecutive batches). We estimated drift separately for each recording segment. Sorting was performed on 5-, 10-, 15- and 45-minute subsets. The match rate and total number of good units sorted are shown as a function of training period (Supplemental Figure 3b-c). In simulated data, match rate was only slightly diminished by the drift rate (regression slope = -0.02, r=-0.51, p<$10^{-3}$) (Supplemental Figure 3d).

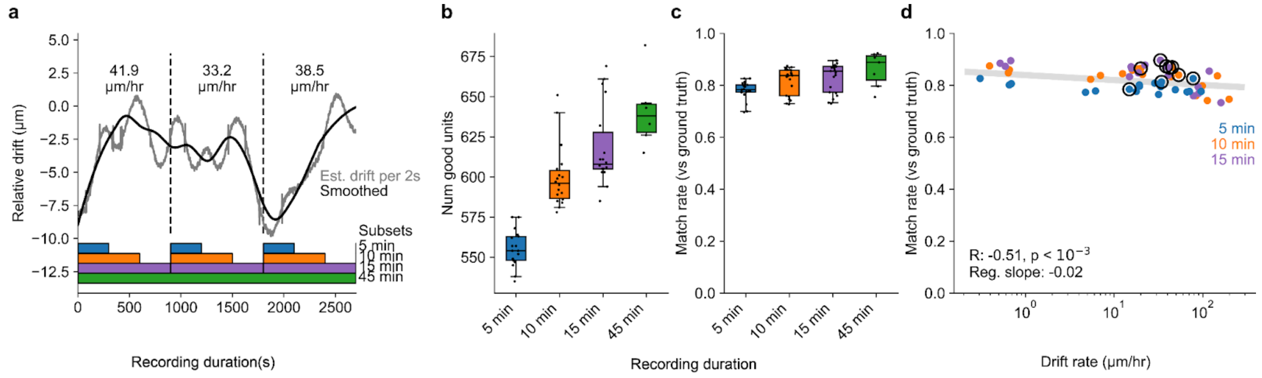


**Supplemental Figure 3. Effect of recording duration and drift on spike sorting in simulated data.** A, Estimated drift rate for example simulated recording with realistic drift. Sorting was performed on 5-, 10-, 15- and 45-minute subsets. B, Number of good units identified by Kilosort4 for each subset. Boxes are

25/75% quartiles, center is median. Whiskers indicate the 95th and 5th percentiles. C, Proportion of 600 ground truth units correctly identified by Kilosort4 (“match rate”) for varying durations of recordings. D, Match rate as a function of drift rate, colored by subset, with linear regression fit. Circled points are examples in a.

Next, using real data, for which ground truth is unavailable, we examined the extent to which we could retrieve units obtained in full recordings using subsets of varying durations (Supplemental Figure 4a-d). To do this, we employed a large, recently published dataset of neurophysiological recordings from macaque prefrontal cortex[21]. This dataset was used because the large number of recordings provided the opportunity to assess the impact of variations in recording stability and unit yield on LSS. We leveraged that variation and assessed the effect of probe drift during the training period on the match rate, as in with the simulated data. Supplemental Figure 4a shows a drift trace along with its estimated drift rates for an example real recording. As with the simulated data, we measured the match rate and total number of good units sorted as a function of training period (Supplemental Figure 4b-c). For the latter, the proportion was computed relative to the sorting result from the full recording. Similar to the simulated data, we observed a negative correlation between match rate and probe drift rate (r=-0.53, $p<10^{7}$)(Supplemental Figure 4d). This result emphasizes the importance of recording stability before establishing the training period for live spike sorting.

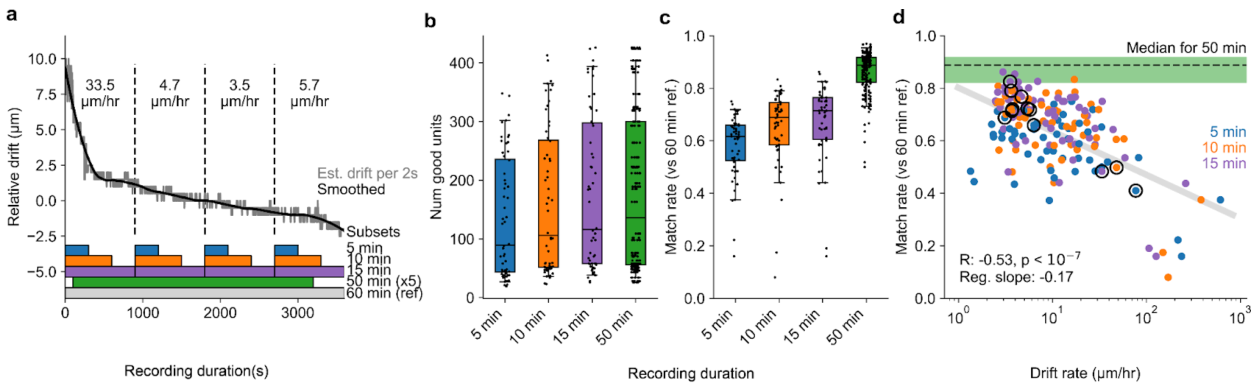

**Supplemental Figure 4. Effect of recording duration and drift on spike sorting in real data.** A, Estimated drift rate for example real recording. Sorting was performed on 5-, 10-, 15- and 45-minute subsets. B, Number of good units identified by Kilosort4 for each subset across several recordings. Boxes are 25/75% quartiles, center is median. Whiskers indicate the 95th and 5th percentiles. C, Proportion of 'good' units correctly identified by Kilosort4 in various durations of recordings, relative to full recording D, Match rate as a function of drift rate, colored by subset, with linear regression fit. Circled points are examples in a.
